# Children with Cerebral Palsy Return to Baseline Community Arm Movement after Constraint Induced Movement Therapy

**DOI:** 10.1101/667246

**Authors:** Brianna M. Goodwin, Emily K. Sabelhaus, Ying-Chun Pan, Kristie F. Bjornson, Kelly L. D. Pham, William O. Walker, Katherine M. Steele

**Affiliations:** Department of Mechanical Engineering, University of Washington, Seattle, WA, USA; Rehabilitation Medicine, Seattle Children’s Hospital, Seattle, WA, USA; Department of Bioengineering, University of Washington, Seattle, WA, USA; Department of Pediatrics, University of Washington, Seattle WA, USA; Physical Medicine & Rehabilitation, University of Washington, Seattle, WA, USA

**Keywords:** Rehabilitation, Wearable Technology, Occupational Therapy, Accelerometer, Upperextremity, Hemiplegia

## Abstract

**Importance:** Constraint Induced Movement Therapy (CIMT) is a common treatment for children with unilateral cerebral palsy (CP). While clinic-based assessments have demonstrated improvements in arm function after CIMT, quantifying if these changes are translated and sustained outside of a clinic setting remains unclear.

**Objective:** Accelerometers were used to quantify arm movement for children with CP one week before, during, and 4+ weeks after CIMT and compared to typically-developing (TD) peers.

**Design:** Observational during CIMT

**Setting:** Clinical assessments and treatment occurred in a tertiary hospital and accelerometry data were collected in the community

**Participants:** 7 children with CP (5m/2f, 7.4 ± 1.2 yrs) and 7 TD peers (2m/5f, 7.0 ± 2.3 yrs)

**Intervention:** 30-hour CIMT protocol

**Outcomes and Measures:** The use ratio, magnitude ratio, and bilateral magnitude were calculated from the accelerometry data. Clinical measures were evaluated before and after CIMT and surveys were used to assess the feasibility of using accelerometers.

**Results:** Before CIMT, children with CP used their paretic arm less than their TD peers. During therapy, their frequency and magnitude of paretic arm use increased in the clinic and in daily life. After therapy, although clinical scores improved, children reverted to baseline accelerometry values. Additionally, children and parents in both cohorts had positive perceptions of wearing accelerometers.

**Conclusions and Relevance:** The lack of sustained improved accelerometry metrics following CIMT suggest therapy gains did not translate to increased movement outside the clinic. Additional therapy may be needed to help the transfer of skills to the community setting.

**What this Article Adds:** This study compares the movement of children with CP undergoing CIMT in the community setting with their typically developing peers. Additional interventions may be needed in combination with or following CIMT to sustain the benefits of the therapy outside of the clinic.

## Introduction

Cerebral palsy (CP) is a non-progressive neurologic disorder of movement and posture that affects approximately 2 of every 1000 children in the United States (Cans, De-la-Cruz, & Mermet, 2008). Constraint-Induced Movement Therapy (CIMT) is one of the most recommended evidence-based treatment for children diagnosed with hemiparesis or unilateral CP (Novak et al., 2013; Sakzewski, Ziviani, & Boyd, 2013). CIMT has been employed as a therapy technique for almost two decades with this population and involves placing the non-paretic arm in a cast for a prescribed period of time with guided therapy to encourage use of the paretic arm (Taub, Ramey, DeLuca, & Echols, 2004). This therapy aims to create unimanual gains, with the goal of skill transfer to bimanual gains outside of the clinical setting.

Although the frequency and duration of treatment interventions can vary, CIMT has consistently demonstrated clinical improvements for children with CP, including increased scores on the Assisting Hand Assessment (AHA), Quality of Upper Extremity Skills test, and parent-reported paretic arm function (Hoare, Imms, Carey, & Wasiak, 2007). Prior research suggests that casting plays an important role in the increase of positive outcomes after CIMT by ‘forcing’ the child to use their paretic arm and causing an increase in treatment intensity (Cope, Forst, Bibis, & Liu, 2008).

While these described gains are important, prior research has also demonstrated a consensus for the use of quantitative measures to monitor movement outside the clinic (Uswatte et al., 2000). Advances in wearable technology have introduced new methods to more easily and accurately track human movement within and outside of the clinic. Specifically, since the 1980’s, improvements in battery life, memory capabilities, cost, and size have made accelerometers an attractive solution for many research applications. Hildebrand & colleagues (2014) reported that accelerometers are the most commonly used objective measure of physical activity and Borghese & colleagues (2017) called accelerometers the gold standard for monitoring physical activity in children.

Accelerometers have been used extensively to monitor steps and physical activity for children with CP and their typically-developing (TD) peers. While accelerometers have been used to monitor arm movement among adult stroke survivors, few studies have used similar methodologies for children with CP (Bailey, Klaesner, & Lang, 2015). Additionally, even fewer studies have investigated how therapy gains in children with CP have transferred learned skills to daily life. Gordon & colleagues (2007) used accelerometry metrics during a standardized clinical test (AHA) before and after CIMT to determine frequency of movement during the assessment. Similarly, Beani & colleagues (2019) used accelerometers to quantify how the hands were being used together in the AHA (Beani et al., 2019). Coker-Bolt & colleagues (2017) used accelerometers to monitor arm movement of children undergoing CIMT. The group reported five out of twelve children receiving CIMT increased frequency of paretic arm movement compared to their non-paretic arm (measured by a use ratio) outside of the clinic after CIMT, although sensors were only worn for six-hours on one day, roughly 1-2 weeks before and after therapy (Coker-Bolt et al., 2017). Due to variability in day-to-day activities, three days of data collections has been reported as necessary to achieve reliable estimates of movement patterns with accelerometers (Mitchell, Ziviani, & Boyd, 2015). Additionally, it is unclear if these increases would be sustained at greater lengths of time following CIMT. To the best of our knowledge, no study has compared the amount of arm movement for children with CP before, during, and after CIMT to that of a group of TD peers, or reported the perspectives of parents and children of wearing accelerometers.

The purpose of this study was to use wearable accelerometers to monitor paretic and non-paretic arm movement of children with CP before, during, and after CIMT and compare their arm movement to TD peers. We hypothesized that during CIMT, children with CP would increase paretic arm frequency and magnitude of use, which would be sustained following CIMT. This study also aimed to examine perceptions of wearing accelerometers as a part of regular clinical care. Examining arm movement outside of the clinic can help to explain the transfer of therapeutic gains from therapy into daily life.

## Methods

### Enrollment and Study Design

Seven children with CP and seven TD children were included in this study (Table 1). The CP cohort was recruited through the CIMT program at a tertiary children’s hospital. Six of the seven participants had prenatal or perinatal strokes/injuries, however one participant had a postnatal stroke at age five affecting the child’s dominant side. Families enrolled in the CIMT program were given information about participation in this study and, if desired, were enrolled. TD children were recruited from the local community. This study was approved by the institutional review board.

**Table 1:** Participant demographics.

| Participant | TD cohort |  |  |  | CP cohort |  |  |  |  |  |
| --- | --- | --- | --- | --- | --- | --- | --- | --- | --- | --- |
|  | Age (years) | Sex | Height (cm) | Weight (kg) | Age (years) | Sex | Height (cm) | Weight (kg) | MACS* | Etiology |
| 1 | 7 | Female | 128 | 26.1 | 7 | Male | 128 | 23.9 | 2 | Prenatal or perinatal stroke |
| 2 | 6 | Female | 124 | 28.3 | 7 | Female | 113 | 18.4 | 2 | Prenatal injury |
| 3 | 2 | Male | 93 | 13.8 | 6 | Male | 119 | 23.8 | 3 | Postnatal stroke |
| 4 | 9 | Female | 139 | 29.8 | 8 | Female | 130 | 32.6 | 3 | Perinatal stroke |
| 5 | 9 | Female | 141 | 26.4 | 7 | Male | 118 | 19.9 | 2 | Perinatal stroke |
| 6 | 7 | Male | 116 | 22.6 | 7 | Male | 139 | 50.0 | 1 | Prenatal or perinatal stroke |
| 7 | 9 | Female | 130 | 25.4 | 10 | Male | 136 | 32.0 | 2 | Perinatal Intraventricular hemorrhage |
| Average (SD) | 7.0 (2.3) | 2 Male<br>5 Female | 124.4 (15.1) | 24.6 (4.9) | 7.4 (1.2) | 5 Male<br>2 Female | 126.1 (9.0) | 28.7 (10.1) | 2.1 (0.7) |  |

\* Manual Ability Classification System

### CIMT Protocol

All participants in the CP cohort received CIMT following the standard protocol at our institution. A custom long-arm, univalve, fiberglass cast was fabricated, which extended from the child’s axillary region to beyond the distal end of the phalanxes. For three weeks, unimanual training of the paretic arm occurred two hours/day, four days/week (Monday-Thursday) and bimanual training two hours/day, once a week (Friday). The cast was worn in and out of therapy throughout the week except for the two-hour period on Friday when bimanual training occurred and the children’s skin was inspected. The training occurred in a small group setting; one therapist to two children with assistance from an adult volunteer. During therapy, the focus was on using shaping techniques to impact the upper extremity; shaping involves using motivating activities of the appropriate difficulty level to allow the child to have successful experiences while developing new skills (DeLuca, Echols, & Ramey, 2007). The goal of shaping is to un-train the ‘developmental disregard’ associated with the paretic arm. This was accomplished when the occupational therapist or occupational therapy assistant gave positive verbal and/or visual recognition to the child when they accomplished challenging tasks. As therapy continued, these positive ques were only given for more complex tasks, with the goal of encouraging the child to perform more challenging movements. Throughout the three-week protocol extensive practice occurred to ensure skills acquisition.

### Wearable Accelerometers

All children were fit with tri-axial ActiGraph GT9X Link accelerometers *(ActiGraph Corp., Pensacola, FL)* that they wore on bilateral wrists. The Actigraphs were placed on each wrist (over cast on paretic arm during CIMT). Parents and child were provided written and verbal instructions on basic wear and use. Data were collected at 100 Hz while the children were awake and not bathing or in water. Data were downloaded through ActiLife *(ActiGraph Corp., Pensacola, FL)* and 1-second epoch activity counts were used for analysis, similar to prior research (Lang, Waddell, Klaesner, & Bland, 2017). Periods that reflected times of non-wear were excluded from data analysis. The CP cohort wore the accelerometers for three days during three periods: (1) one week before CIMT, (2) during the second week of CIMT, and (3) 4+ weeks after CIMT. The periods before and after CIMT were designed to align with clinical exams and varied slightly for each participant due to scheduling and family commitments. On average, children were seen 7 ± 2 days prior to CIMT and 7.6 ± 3.9 weeks after completion of CIMT. Five of the seven participants were seen 4-6 weeks following CIMT, and due to time constraints the other two participants were seen at weeks 11 and 15. The TD cohort also wore the accelerometers for three, three-day periods temporally spaced to align with the CIMT protocol, but no intervention occurred. For all outcome measures, averages across the three periods were used for the TD cohort.

To evaluate magnitude and amount of arm movements, three outcome metrics were used: use ratio, magnitude ratio, and bilateral magnitude (Bailey et al., 2015; Urbin, Waddell, & Lang, 2015). The paretic and non-paretic arm will be referred to as the non-dominant and dominant arm, respectively, while describing the metrics used. Use ratio provides a measurement comparing the frequency of activity between the right and left arm:

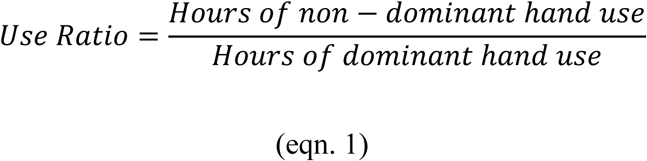

A use ratio equal to one indicates that the non-dominant and dominant arms were used an equal amount of time throughout the day, while values greater than one would indicate greater use of the non-dominant arm. To calculate hours of arm movement, the number of epochs with activity counts greater than zero were summed and converted to hours for each arm. Use ratio is often used since it gives a single numerical output from a large amount of data, which can be used to compare the activity frequency of each arm (Urbin et al., 2015).

The magnitude ratio was calculated as a metric to compare the magnitude of acceleration of the arms at each time point:

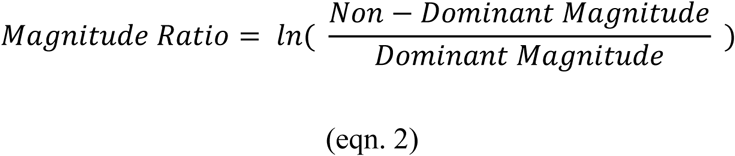

The magnitude was calculated by taking the vector magnitude of the activity counts for each epoch. Similar to prior research, the natural log was used to avoid skewness in the ratio caused by an underestimation of the denominator (van der Pas, Verbunt, Breukelaar, van Woerden, & Seelen, 2011). Values greater than 7 or less than -7 were set to 7 and -7, respectively. The average magnitude was calculated for each period. A magnitude ratio near zero indicates similar use of each arm, while a negative number indicates more dominant arm use, and a positive number indicated more non-dominant arm use.

The bilateral magnitude was used to compare the overall movement of both arms together as a measure of bilateral arm movement:

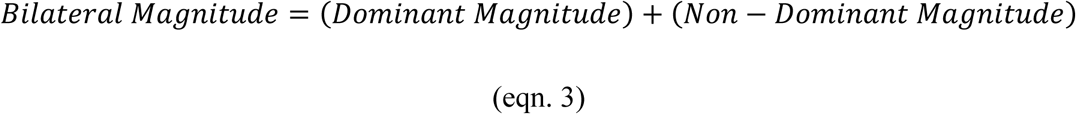

Similar to the magnitude ratio, the vector magnitude of the activity counts was calculated for each epoch. A greater bilateral magnitude indicates greater overall movement of both arms.

### Clinical Measures

Before and after CIMT, the occupational therapist performed a comprehensive evaluation for each child with CP. The grip, pinch, and lateral pinch of the child’s paretic and non-paretic arms were measured using pinch and grip dynamometers; the maximum score over three trials was recorded. The Canadian Occupational Performance Measure (COPM), a client-centered clinical practice outcome assessment, was used to measure patient identified problems of daily function (Law et al., 1990). Children identified goals of self-care, productivity, or leisure and scored themselves on their performance and satisfaction. Finally, the Box & Blocks Test was used as a standardized measure of coordination.

### Survey Data

Parents of both cohorts were given surveys after the second (during CIMT for the CP cohort) and third (after CIMT for the CP cohort) data collection to assess parent and child attitudes toward wearing wearable sensors as a part of clinical care. Seven questions asked parents to indicate their level of agreeance with statements aimed at understanding the benefits and challenges families experienced in using wrist-worn accelerometers. The questions assessed comfort, aesthetics, and parents’ interest in accessing their child’s personalized accelerometry data. The survey also included three open-ended questions for parents to provide specific feedback regarding how to improve the technology for their child.

### Statistical Analysis

Due to the study’s small sample size, nonparametric tests were used for all comparisons. Friedman’s Test were used to compare between visits for each cohort and Wilcoxon Rank-Sum Tests were used to evaluate differences between cohorts or time points (α=0.05). All data analysis and statistical tests were conducted using custom programs in MATLAB (*MathWorks, Inc., Natick, MA)*.

## Results

Before therapy, children with CP used their paretic arm significantly less than their non-paretic arm or their TD peers in daily life. The average use ratio of the CP cohort before therapy was 0.79 ± 0.03 versus 0.96 ± 0.03 for the TD cohort (p = 0.026, Figure 1). The magnitude ratio of the CP cohort during this pre-therapy period was significantly less than the TD cohort at -1.70 ± 0.28, compared with -0.28 ± 0.24 for the TD cohort (p = 0.026). Even with the decrease in paretic arm movement, the combined arm movement of the CP cohort during this period stayed similar to the TD cohort; the bilateral magnitude was 111.6 ± 24.3 for the CP cohort and 126.7 ± 27.4 for the TD cohort (p = 0.46). There was no significant difference in TD arm movement across the three time periods for the use ratio (p = 0.56), magnitude ratio (p = 0.28), or bilateral magnitude (p = 0.066).

**Figure 1:**
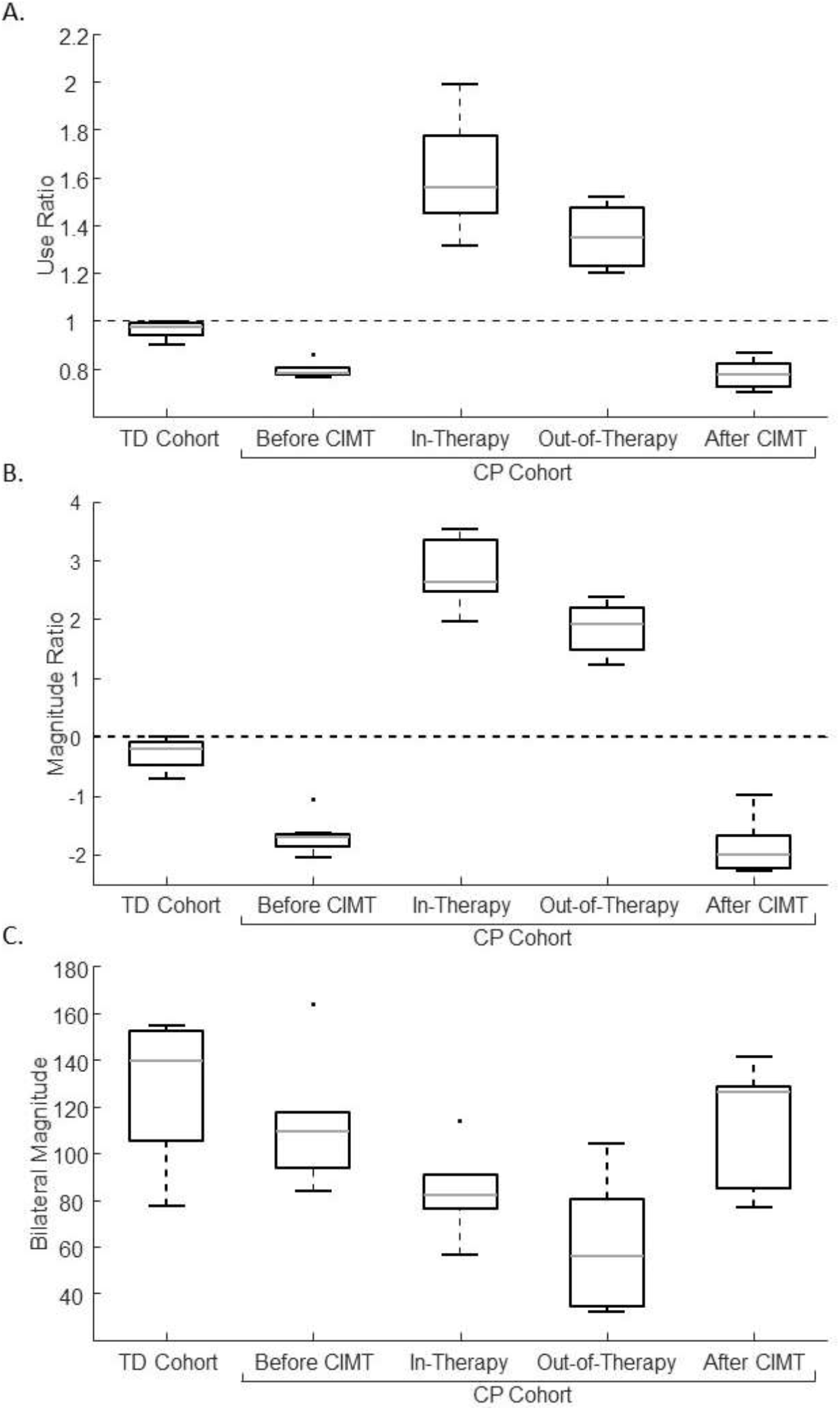
Use ratio (A), magnitude ratio (B), and bilateral magnitude (C) for the TD and CP cohorts. The averages over the three time periods are shown for the TD cohort. For the CP cohort, results are shown for each period: before CIMT, during CIMT (including both time in-therapy at the hospital and out-of-therapy), and after CIMT. The dashed lines in A and B show equal arm movement – a value of one for use ratio and zero for magnitude ratio.

During CIMT, the children with CP significantly increased their use ratio and magnitude ratio both in-therapy (while working with a therapist) and out-of-therapy (at school, home, etc.) compared to before CIMT (Figure 1). The use ratio increased to 1.61 ± 0.21 (p = 0.00058) while in-therapy and 1.36 ± 0.12 (p = 0.0023) while out-of-therapy, but still wearing the cast.

Similarly, the magnitude ratio increased to 2.80 ± 0.53 (p = 0.00058) in-therapy and 1.85 ± 0.40 (p = 0.0023) out-of-therapy. There was a significant difference in the use ratio and magnitude ratio when comparing the time in-therapy and time out-of-therapy (p = 0.026 and p = 0.0041, respectively). Participant’s overall movement, measured by the bilateral magnitude decreased both in therapy (84.19 ± 16.12, p = 0.018) and out of therapy (58.95 ± 25.67, p=0.007), suggesting less overall movement of both arms while participating in CIMT.

After CIMT, the CP cohort returned to baseline values for all accelerometry outcomes. The use ratio (0.78 ± 0.06, p = 0.26), magnitude ratio (−1.87 ± 0.42, p = 0.26), and bilateral magnitude (110.0 ± 24.7, p = 0.90) were not significantly different from pre-CIMT values (Figure 1). However, the children with CP had improvements in the clinical measures after CIMT compared to pre-CIMT (Table 2). There was a significant increase in grip strength and increases, although not significant, in 3-point pinch, lateral grasp, and Box & Blocks Test) for the paretic arm. Additionally, the children with CP ranked themselves as more able to reach their self-identified goals following CIMT, as measured by the COPM. The children rated themselves higher on their performance and satisfaction with reaching their bimanual goals and unimanual goals after CIMT.

**Table 2:** Change in clinical measures after CIMT.

|  | Paretic | Non-Paretic |
| --- | --- | --- |
| Grip (lbs) | 8.3 ± 7.3<br>(p = 0.05) | -0.1 ± 1.3<br>(p = 0.89) |
| 3-Point pinch (lbs) | 3.2 ± 5.0<br>(p = 0.40) | 0.25 ± 0.25<br>(p = 1.0) |
| Lateral grasp (lbs) | 1.5 ± 1.0<br>(p = 0.12) | -1.3 ± 0.1<br>(p = 0.10) |
| Box & Blocks (blocks) | 4.3 ± 5.1<br>(p = 0.38) | -0.3 ± 5.9<br>(p = 0.97) |
| COMP unimanual (performance/satisfaction) | 2.2 ± 1.1/2.5 ± 1.3<br>(p = 0.009/p = 0.007) |  |
| COMP bimanual (performance/satisfaction) | 4.6 ± 2.5/4.8 ± 2.6<br>(p = 0.0003/p = 0.0003) |  |

Families in the both the TD and CP cohorts had positive perceptions of wearing accelerometers, but were generally neutral on whether or not accelerometers should be integrated into clinical care (Table 3). Families in both cohorts emphasized the comfort of the sensors and wanted to learn from the data gained from the sensors, specifically about how their child’s upper extremity movement and function had changed following CIMT. Additionally, parents suggested that their child would have been more compliant wearing the sensors if the sensors interacted with the child using lights, for example. There were no statistical differences between cohorts in questionnaire responses.

**Table 3:** Parent survey responses (1 = strongly disagree, 5 = strongly agree)

|  | CP | TD |
| --- | --- | --- |
| Sensors were comfortable for my child to wear | 4.2 ± 0.8 | 4.0 ± 0.7 |
| My child did not want to wear the sensors | 1.7 ± 0.9 | 2.2 ± 0.8 |
| My child enjoyed wearing the sensors | 4.0 ± 0.7 | 3.6 ± 0.8 |
| Sensors interfered with daily activities | 2.3 ± 1.0 | 1.7 ± 0.9 |
| The sensors were a conversation starter | 4.3 ± 0.6 | 4.3 ± 0.5 |
| I wish we could learn more about the data from the sensors | 4.2 ± 0.6 | 3.9 ± 1.1 |
| Wearable sensors should be part of clinical care | 3.6 ± 0.9 | 3.4 ± 1.0 |

## Discussion

CIMT is a common therapy for children with unilateral CP, which aims to achieve unimanual gains that will transfer to bimanual arm use outside of the clinic and be sustained after the completion of the therapy. The purpose of this study was to quantify arm use before, during, and after CIMT in comparison to TD peers using accelerometry. Among a cohort of seven children with unilateral CP, we found that the therapy combined with the cast increased the amount (use ratio) and intensity (magnitude ratio) of paretic arm use, both while actively engaged in therapy at the hospital (2 hrs/day), but also in the community, confirming that the cast effectively increases use and practice of the paretic arm in both environments. However, results demonstrated a decrease in overall activity level during CIMT (bilateral magnitude), especially during time periods outside of therapy, which may suggest that casting has a detrimental effect on activity and participation in home and community activities. Further, following CIMT there was a return to baseline for all accelerometer measures, indicating that gains in paretic arm use were not maintained after therapy concluded. These results suggest that new strategies or home exercise programs, such as remind-to-move or other follow-up programs may be necessary to maintain increased paretic arm use and improved functional skills in daily life following CIMT (A.-Q. V. Dong & Fong, 2016).

Accelerometers have been previously used in children with CP to evaluate movement (Sokal, Uswatte, Vogtle, Byrom, & Barman, 2015) and our results parallel many prior findings. Focused on CIMT, Coker-Bolt & colleagues (2017) used accelerometers to evaluate arm movement for one day before and after a week-long CIMT camp. While they did observe improvements in five of the twelve participants 1-2 weeks following CIMT, we did not see similar improvements 4+ weeks after the three week-long intervention. There may indicate immediate gains in arm movement following CIMT for some children, however additional intervention may be needed to maintain these improvements. The use ratio and magnitude ratios reported here had a much smaller range than those reported by Coker-Bolt & colleagues (2017). This may be due to the fact that they obtained only one day of data collection, while our study had children wear the accelerometers for three days during each of the three data collection periods. This could also be due to a difference in impairment severity, as some of the baseline use ratio and magnitude ratio values reported by Coker-Bolt & colleagues (2007) were closer to our TD cohort. The differences in magnitude ratio between cohorts was similar to that of the asymmetry index presented by Beani & colleges (2019), demonstrating that, in both studies, children with CP used their paretic arm significantly less than TD peers.

The use of accelerometry data also allowed comparison with common clinical tests that are standard of care for children with CP. Similar to prior research, clinical measures improved after CIMT including improvements in grip and pinch strength (Martin, Burtner, Poole, & Phillips, 2008) and patient or parent reported goals (Gordon, 2011). In examining COPM, 63% of the goals chosen by the children with CP cohort in this study were bimanual goals, similar to 85% previously reported by Gordon & colleagues (2011). While CIMT emphasizes unilateral practice of the paretic-arm, these patient-reported goals and decreases in bilateral magnitude found in this study may support combinations of CIMT with bimanual therapy. Additionally, a remind to move (RTM) protocol has been used in other studies to create more self-awareness of the paretic arm (A.-Q. V. Dong & Fong, 2016; V. A. Dong, Fong, Chen, Tseng, & Wong, 2017). The RTM protocol involves wearing a sensory cuing device on the paretic arm that vibrates every 15 minutes and reminds the child to use their paretic arm. To the best of our knowledge, no study has compared the effects of traditional CIMT to CIMT followed by RTM, however the combination could theoretically facilitate the transfer of unimanual skills gained in CIMT into bimanual tasks in daily living.

Results of this study are limited based upon the small number of children in each cohort. Because CP encompasses such a heterogeneous population, there may not be enough children in this cohort to represent trends for the entire CP population. Furthermore, because of the small sample size, conclusions regarding which children benefit most from this therapy cannot be drawn. Independent factors including age, onset of hemiplegia, location of brain injury, or side of hemiplegia may influence the benefits of CIMT, but these trends cannot be determined from the current data set. Additionally, there are numerous variations of CIMT with different frequencies, intensities, and total durations (Sakzewski, Provan, Ziviani, & Boyd, 2015). The data and analysis outcomes presented here are only applicable for this specific protocol, with a total dosage time of 30 hours. The accelerometry results described in this report were recorded from wrist-worn accelerometers; giving only overall arm movements. This analysis assumed if a child improved their finger movements they also would have improved their arm movements; at this time we do have the technology to measure finger movements accurately outside of the clinic.

### Implications for Occupational Therapy Practice and Research

The results of this study have the following implications for occupational therapy practice and research:

- A 30-hour CIMT protocol results in increased grip strength of the paretic hand and improvement of a child’s perception of their ability to accomplish his/her goals.
- During therapy, wearing the cast in and out of therapy promoted more paretic arm movement compared to before CIMT. However, there was an overall decrease in movement of both arms while wearing this cast which should be taken into consideration when developing home activities during CIMT.
- Following CIMT, children returned to baseline values for arm movement in daily life, suggesting that additional strategies may be needed to translate gains from CIMT into activities of daily living in their home, school, and community settings following CIMT.
- Families and children in both cohorts had positive perceptions and experiences using accelerometers as a method to monitor movement outside of the clinic.

## Conclusion

To evaluate the benefits of a 30-hour CIMT protocol, accelerometers were worn on both wrists by children with CP for three-day data collection periods before, during, and after therapy. Our results suggest that, while the CP cohort improved accelerometry-measured metrics of arm movement during therapy, all metrics fell back to baseline values after therapy. However, children did show improvements in standard measures of clinical function. The lack of sustained accelerometry improvements following CIMT suggests unimanual skills gained in therapy are not be maintained 4+ weeks following therapy. Parents and children had positive perceptions of wearing accelerometers which support their further use to monitor movement and inform care for children with CP.

## Acknowledgements

This project was supported by the Seattle Children’s Hospital Academic Enrichment Fund and NIH NIBIB R01-EB021935. We would like to thank the occupational therapists, assistants, and families for their time and engagement, without whom this research would not be possible.

## References

Bailey, R. R., Klaesner, J. W., & Lang, C. E. (2015). Quantifying real-world upper-limb activity in nondisabled adults and adults with chronic stroke. Neurorehabilitation and neural repair, 29(10), 969–978.

Beani, E., Maselli, M., Sicola, E., Perazza, S., Cecchi, F., Dario, P., … Sgandurra, G. (2019). Actigraph assessment for measuring upper limb activity in unilateral cerebral palsy. Journal of neuroengineering and rehabilitation, 16(1), 30.

Borghese, M., Tremblay, M., Leblanc, A., Leduc, G., Boyer, C., & Chaput, J. (2017). Comparison of ActiGraph GT3X+ and Actical accelerometer data in 9–11-year-old Canadian children. Journal of sports sciences, 35(6), 517–524.

Cans, C., De-la-Cruz, J., & Mermet, M.-A. (2008). Epidemiology of cerebral palsy. Paediatrics and child health, 18(9), 393–398.

Coker-Bolt, P., Downey, R. J., Connolly, J., Hoover, R., Shelton, D., & Seo, N. J. (2017). Exploring the feasibility and use of accelerometers before, during, and after a camp-based CIMT program for children with cerebral palsy. Journal of pediatric rehabilitation medicine, 10(1), 27–36.

Cope, S. M., Forst, H. C., Bibis, D., & Liu, X.-C. (2008). Modified Constraint-Induced Movement Therapy for a 12-Month-Old Child with Hemiplegia: A Case Report. American Journal of Occupational Therapy, 62(4), 430–437.

DeLuca, S., Echols, K., & Ramey, S. L. (2007). ACQUIREc therapy: A training manual for effective application of pediatric constraint-induced movement therapy: Mindnurture.

Dong, A.-Q. V., & Fong, N.-K. K. (2016). Remind to move–A novel treatment on hemiplegic arm functions in children with unilateral cerebral palsy: A randomized cross-over study. Developmental neurorehabilitation, 19(5), 275–283.

Dong, V. A., Fong, K. N., Chen, Y. F., Tseng, S. S., & Wong, L. (2017). ‘Remind-to-move’treatment versus constraint-induced movement therapy for children with hemiplegic cerebral palsy: a randomized controlled trial. Developmental Medicine & Child Neurology, 59(2), 160–167.

Gordon, A. M. (2011). To constrain or not to constrain, and other stories of intensive upper extremity training for children with unilateral cerebral palsy. Developmental Medicine & Child Neurology, 53(4), 56–61.

Gordon, A. M., Schneider, J. A., Chinnan, A., & Charles, J. R. (2007). Efficacy of a hand–arm bimanual intensive therapy (HABIT) in children with hemiplegic cerebral palsy: a randomized control trial. Developmental Medicine & Child Neurology, 49(11), 830–838.

Hildebrand, M., Van, V. H., Hansen, B. H., & Ekelund, U. (2014). Age group comparability of raw accelerometer output from wrist-and hip-worn monitors. Medicine and science in sports and exercise, 46(9), 1816–1824.

Hoare, B., Imms, C., Carey, L., & Wasiak, J. (2007). Constraint-induced movement therapy in the treatment of the upper limb in children with hemiplegic cerebral palsy: a Cochrane systematic review. Clinical Rehabilitation, 21(8), 675–685.

Lang, C. E., Waddell, K. J., Klaesner, J. W., & Bland, M. D. (2017). A method for quantifying upper limb performance in daily life using accelerometers. Journal of visualized experiments: JoVE(122).

Law, M., Baptiste, S., McColl, M., Opzoomer, A., Polatajko, H., & Pollock, N. (1990). The Canadian occupational performance measure: an outcome measure for occupational therapy. Canadian Journal of Occupational Therapy, 57(2), 82–87.

Martin, A., Burtner, P. A., Poole, J., & Phillips, J. (2008). Case report: ICF-level changes in a preschooler after constraint-induced movement therapy. American Journal of Occupational Therapy, 62(3), 282–288.

Mitchell, L. E., Ziviani, J., & Boyd, R. N. (2015). Variability in measuring physical activity in children with cerebral palsy. Medicine and science in sports and exercise, 47(1), 194–200.

Novak, I., Mcintyre, S., Morgan, C., Campbell, L., Dark, L., Morton, N., … Goldsmith, S. (2013). A systematic review of interventions for children with cerebral palsy: state of the evidence. Developmental Medicine & Child Neurology, 55(10), 885–910.

Sakzewski, L., Provan, K., Ziviani, J., & Boyd, R. N. (2015). Comparison of dosage of intensive upper limb therapy for children with unilateral cerebral palsy: How big should the therapy pill be? Research in developmental disabilities, 37, 9–16.

Sakzewski, L., Ziviani, J., & Boyd, R. N. (2013). Efficacy of upper limb therapies for unilateral cerebral palsy: a meta-analysis. Pediatrics, peds. 2013–0675.

Sokal, B., Uswatte, G., Vogtle, L., Byrom, E., & Barman, J. (2015). Everyday movement and use of the arms: Relationship in children with hemiparesis differs from adults. Journal of pediatric rehabilitation medicine, 8(3), 197–206.

Taub, E., Ramey, S. L., DeLuca, S., & Echols, K. (2004). Efficacy of constraint-induced movement therapy for children with cerebral palsy with asymmetric motor impairment. Pediatrics, 113(2), 305–312.

Urbin, M., Waddell, K. J., & Lang, C. E. (2015). Acceleration metrics are responsive to change in upper extremity function of stroke survivors. Archives of physical medicine and rehabilitation, 96(5), 854–861.

Uswatte, G., Miltner, W. H., Foo, B., Varma, M., Moran, S., & Taub, E. (2000). Objective measurement of functional upper-extremity movement using accelerometer recordings transformed with a threshold filter. Stroke, 31(3), 662–667.

van der Pas, S. C., Verbunt, J. A., Breukelaar, D. E., van Woerden, R., & Seelen, H. A. (2011). Assessment of arm activity using triaxial accelerometry in patients with a stroke. Archives of physical medicine and rehabilitation, 92(9), 1437–1442.

